# We the Hunted

**DOI:** 10.1101/2022.09.29.510060

**Authors:** Jesse M. Martin, A.B. Leece, Andy I.R. Herries, Stephanie E. Baker, David S. Strait

## Abstract

Classic depictions of human evolutionary ecology cast *Homo* as predator and other hominins, including *Paranthropus robustus,* as prey. Such hypotheses rest on a small number of fossils that exhibit evidence of carnivore predation, including the iconic SK 54 cranium from Swartkrans in South Africa. Here we demonstrate that the SK 54 cranium shares its closest affinities with *H. erectus* rather than *P. robustus.* Demonstrating that *Homo* was prey for leopards at Swartkrans weakens the historically significant hypothesis that *Homo* was better able to avoid predation because of being behaviourally and technologically advanced compared to *Paranthropus*. Subsequent ideas about hominin paleobiology derived from this hypothesis warrant reconsideration.

## Introduction

SK 54 is a partial hominin cranium that was recovered from the ~1.9-1.8 Ma palaeocave deposits of Swartkrans Member 1 Hanging Remnant in South Africa in 1949 and subsequently prepared by John T. Robinson (Brain 1970; 1981; Herries 2022). This specimen is an iconic hominin fossil that has influenced both the development of the discipline of cave taphonomy as well as narratives concerning how multiple hominin species shared the landscape of Pleistocene southern Africa (Brain 1970). Brain (1970; 1981) described two carnivore puncture marks on its left and right parietal bones (Figure 1) whose location, size, and spacing indicate strongly that they were inflicted by a leopard (famously, the marks conform well to the canines of leopard fossil SK 349 from the same deposit) (Brain 1969). Leopards are known to be predators rather than scavengers (Brain 1970; Hayward et al., 2006) and thus this specimen preserves direct evidence of the predation of hominins. For this reason, although there are other hominin fossils from Swartkrans that exhibit carnivore modification marks (Brain 1970; Brain 1981; White 1988; Pickering 2001), no other specimen has figured as centrally in hypotheses concerning carnivore predation on hominins than SK 54 (Brain 1970; 1981; Holloway 1972; Blumenberg and Todd 1974; Susman 1987; White 1988; Lee-Thorp et al. 2000; Pickering et al. 2004; Pickering 2005; Gommery et al. 2007; Pobiner 2008; Val et al. 2014; Arriaza et al. 2021). The specimen has previously been attributed to *Paranthropus robustus* (Brain, 1970) and that taxonomy has remained unchallenged and current (e.g., Wolpoff 1971; 1974; Kimbel et al. 1984; Rak et al. 2021). This attribution has underwritten hypotheses that australopiths were prey while early *Homo* were transforming into predators, as elucidated in Brain’s (1981) classic monograph, *The Hunters or The Hunted.* Here we provide taxonomic evidence that challenges this narrative. Our analysis of SK 54 results principally from morphological comparisons conducted in South Africa in 2018, 2019 and 2022 on original fossil specimens of *Australopithecus africanus, A. sediba, P. robustus* and early *Homo* curated at the Evolutionary Studies Institute of the University of the Witwatersrand, and the Ditsong Museum of Natural History.

**Figure 1:**
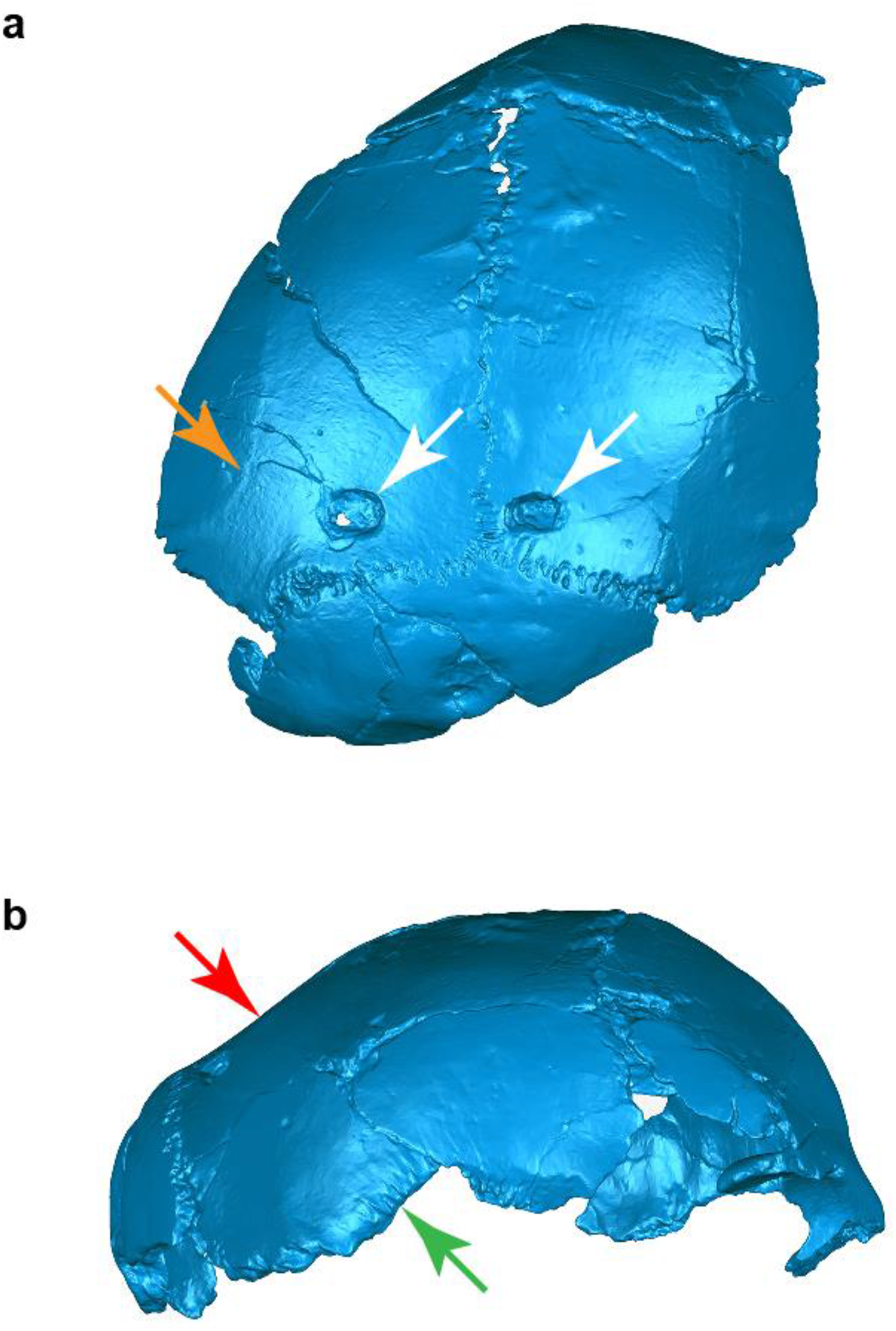
Three-dimensional surface scans of SK 54 shown in **a,** superoposterior and **b,** lateral views. White arrows indicate puncture marks. Orange arrow indicates the left superior temporal line positioned laterally far from the sagittal suture. Red arrow indicates likely pre-lambdoidal flattening. Green arrow indicates the posterior aspect of the parietal portion of the squamosal suture that is straight and shows minimal overlap with the temporal bone.

## Results

SK 54 is a fragmentary neurocranium preserving parts of the occipital, frontal, and left and right parietal bones. The degree of sutural fusion suggests that the specimen may have been a juvenile at the time of death. The specimen is plasticly deformed such that it is not possible to assess overall neurocranial shape, and this deformation precludes meaningful quantitative analysis. Moreover, its neurocranial vault is distorted on both the right and left sides, making a digital reconstruction highly subjective and of little diagnostic value. The individual evidently had a generally small brain, however, estimates of its cranial capacity cannot be made with confidence (e.g., Holloway, Broadfield and Yuan 2003) Nonetheless, aspects of preserved morphology challenge its traditional taxonomic attribution to *Paranthropus* and suggest affinities with *Homo.* The temporal lines are well separated and laterally positioned (Figure 1), a configuration that is incompatible with adult *P. robustus* specimens that exhibit either a sagittal crest (in putative males) or nearly convergent temporal lines (in females) (Lockwood et al. 2007). There are no juvenile *P. robustus* specimens that preserve the requisite morphology to assess this trait but the juvenile *P. boisei* specimen L338y-6 (whose sutures are as or more open than those of SK 54, implying a coarse similarity in age) exhibits well-developed, strongly convergent temporal lines (Rak and Howell 1978), suggesting that SK 54 may not be *Paranthropus.* Kimbel, White, and Johanson (1984) have previously argued that SK 54 preserves rugose *striae parietalis* that they suggest are correlated with a high degree of overlap between the temporal squama and parietal at the squamosal suture. They based their inference on the observation by Rak (1978) that juvenile *H. sapiens* from a Holocene population exhibited fine rather than rugose striae. However, the length of the *striae parietalis* preserved in SK 54 (Figure 1) are notably less than those of L338y-6 and adult *P. robustus* specimens DNH7, DNH152, and DNH155, and both the length and rugosity of SK 54’s striae closely resemble the condition in the juvenile *H. erectus sensu lato* specimen DNH 134. In our assessment, enough of the inferior bevelled edge of the right parietal (Figure 1) is preserved to indicate that the temporal and parietal portions of the squamosal suture would not have overlapped extensively, unlike the extensive overlap seen in both adult and juvenile *Paranthropus* (e.g., Rak 1978; Rak and Howell 1978; Herries et al. 2020). Finally, *contra* Kimbel, White, and Johanson (1984), the very wide separation of the superior temporal lines on the frontal bone argues against the inference that a frontal trigon would have developed in adulthood.

Two discrete traits of SK 54 may suggest affinities with *Homo erectus sensu lato.* The preserved posterior portion of the squamosal suture is straight as it rises anteriorly and superiorly (Figure 1). Moreover, SK 54’s parietal bones are flattened anterior to the lambdoidal suture (Figure 1), although there is distortion present in this region. Pre-lambdoidal flattening is a derived characteristic of many *H. erectus* specimens. The fragmentary and deformed nature of SK 54 precludes a definitive taxonomic allocation but, heuristically, superimposing SK 54 onto *H. erectus sensu lato* specimen KNM-ER 42700 (Spoor et al., 2007) demonstrates a striking similarity between the two specimens (Figure 2). Minimally, a provisional assignment of SK 54 to *Homo* seems warranted and a tentative species-level allocation to *H. erectus sensu lato* is plausible. This assignment adds to the evidence for *Homo* at Swartkrans Member 1 Hanging Remnant that includes the juvenile cranium SK 27 that Clarke (1977) reclassified from *P. robustus* to *Homo,* and the suggested partial skull consisting of individual fossils SK 80, SK 846b, SK 847 (Clarke, Howell, & Brain 1970) and sometimes also the mandible fragment SK 45 (Brain 1981). At 1.9-1.8 Ma (Herries 2022), these fossils are slightly younger than the 2.04-1.95 Ma DNH 134 cranium from Drimolen Main Quarry that also shows affinities to *Homo erectus* (Herries, Martin, & Leece et al., 2020).

**Figure 2:**
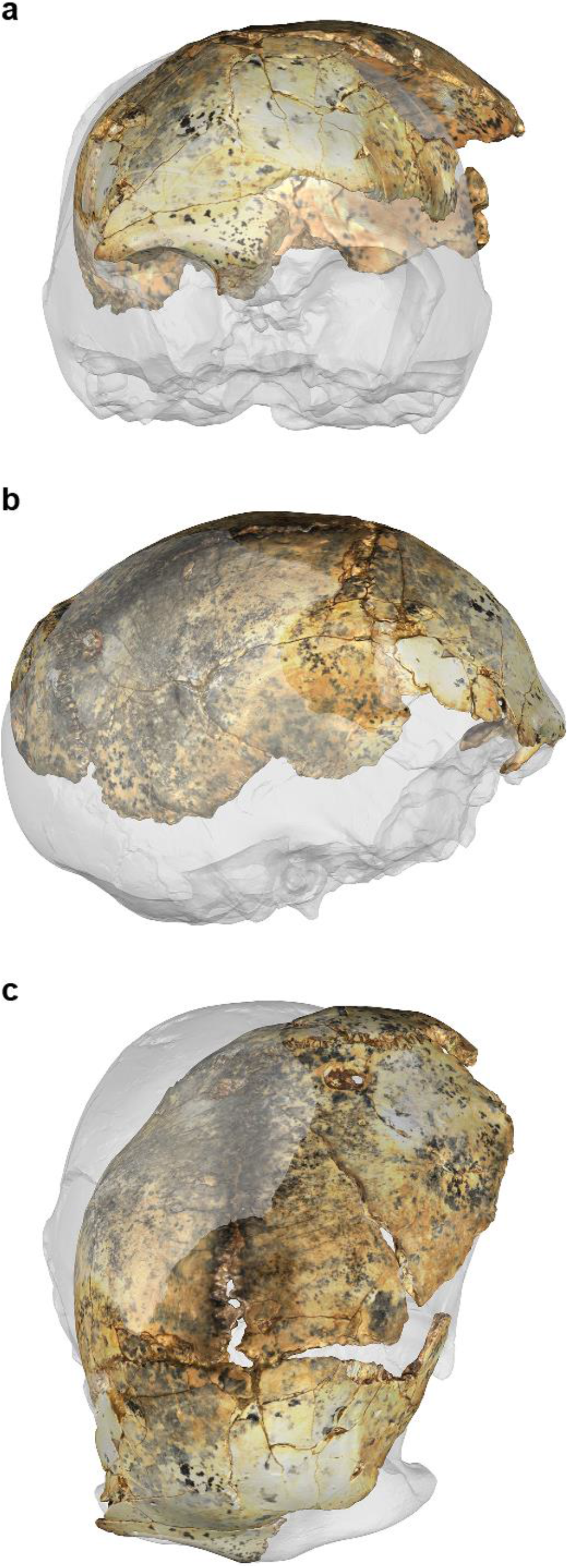
Superimposition of a three dimensional surface scan of SK 54 onto a cast of KNM-ER 42700 (transparent, scaled to 90% of its size) in **a,** frontal, **b,** right lateral, and **c,** superior views. The specimens are aligned at their right orbital margins and circumorbital regions, which are the least distorted portions of SK 54.

## Discussion

Brain’s work on cave taphonomy remains seminal, and his taphonomic assessment linking SK 54 with leopard predation is unchallenged. Our taxonomic reassessment of SK 54 demonstrates that *Homo* was also prey for leopards in the early Pleistocene, and this characterisation could not be further removed from classic depictions of *Homo* the hunter and *Paranthropus* the hunted. Allocating SK 54 to *Homo* tempers the impetus for supposing that early *Homo* and *P. robustus* were differentially predated because of the former’s behavioural and technological advancement.

It is impossible to know with certainty how the history of palaeoanthropology might have been different had SK 54 been recognized as *Homo w*hen it was first discovered, but it is reasonable to infer that the impact of such a realization would have been significant. Only six years prior to the publication of Brain’s (1970) now-classic paper, the description of the newly discovered *H. habilis* (Leakey et al. 1964) included an assessment of the relative tool-making skills and trophic positions of the new species and its contemporary, *Zinjanthropus boisei* (i.e., *P. boisei*), “While it is possible that *Zinjanthropus* and *Homo habilis* both made stone tools, it is probable that the latter was the more advanced tool maker and that the *Zinjanthropus* skull represents an intruder (or a victim) on a *Homo habilis* living site.” Shortly thereafter, the highly influential *Man the Hunter* conference was held followed by the publication of its accompanying edited volume (Lee & DeVore 1968; Barr et al. 2020) that described hunting as a fundamentally important human adaptation. Brain’s (1970; 1981) interpretation of SK 54, based on an incorrect taxonomy was therefore compatible with the thinking of the time but it could have instead been a powerful challenge to conventional wisdom. Ideas, like species, evolve and have descendants, so the evidence presented here should prompt a reassessment of hypotheses concerning the biology, behaviour, and technological capabilities of *Homo* and *Paranthropus* that are derived from earlier ideas positing *Homo* as predator, and *Paranthropus* as prey (for example, Lockwood et al. 2007). Our findings complement a recent zooarchaeological analysis showing that the appearance of *H. erectus* is not associated with increased evidence for hominin carnivory (Barr et al. 2020).

